# The pomegranate (*Punica granatum* L.) genome provides insights into fruit quality and ovule developmental biology

**DOI:** 10.1101/158857

**Authors:** Zhaohe Yuan, Yanming Fang, Taikui Zhang, Zhangjun Fei, Fengming Han, Cuiyu Liu, Min Liu, Wei Xiao, Wenjing Zhang, Mengwei Zhang, Youhui Ju, Huili Xu, He Dai, Yujun Liu, Yanhui Chen, Lili Wang, Jianqing Zhou, Dian Guan, Ming Yan, Yanhua Xia, Xianbin Huang, Dongyuan Liu, Hongmin Wei, Hongkun Zheng

## Abstract

Pomegranate *(Punica granatum* L.) with an uncertain taxonomic status has an ancient cultivation history, and has become an emerging fruit due to its attractive features such as the bright red appearance and the high abundance of medicinally valuable ellagitannin-based compounds in its peel and aril. However, the absence of genomic resources has restricted further elucidating genetics and evolution of these interesting traits. Here we report a 274-Mb high-quality draft pomegranate genome sequence, which covers approximately 81.5% of the estimated 336 Mb genome, consists of 2,177 scaffolds with an N50 size of 1.7 Mb, and contains 30,903 genes. Phylogenomic analysis supported that pomegranate belongs to the Lythraceae family rather than the monogeneric Punicaceae family, and comparative analyses showed that pomegranate and *Eucalyptus grandis* shares the paleotetraploidy event. Integrated genomic and transcriptomic analyses provided insights into the molecular mechanisms underlying the biosynthesis of ellagitannin-based compounds, the color formation in both peels and arils during pomegranate fruit development, and the unique ovule development processes that are characteristic of pomegranate. This genome sequence represents the first reference in Lythraceae, providing an important resource to expand our understanding of some unique biological processes and to facilitate both comparative biology studies and crop breeding.

## Introduction

Pomegranate *(Punica granatum* L.), which is native to central Asia, is an ancient medicinal fruit crop that is grown worldwide[1] and has considerable economic value. Compared to other fruit crops, such as orange *(Citrus sinensis),* apple *(Malus domestica),* grape *(Vitis vinifera),* and kiwifruit *(Actinidia chinensis),* pomegranate has higher levels of antioxidants (~11.33 mmol/100 g)[2], which are potentially beneficial in preventing cardiovascular disease, diabetes, and prostate cancer[3]. Consequently, pomegranate is also referred as a ‘super fruit’[4] and the planted acreages and fruit production of pomegranate have been increased substantially over the past decade[5].

Although the genus *Punica* was previously placed in its own monogeneric family (Punicaceae), recent morphological[6] and molecular[7] evidence, as well as the new classification in the APG IV system[8], suggest that it is a member of Lythraceae. Pomegranate has become an attractive system for studying several valuable biological features, such as the metabolism of ellagitannin-based compounds[9], color formation in the fruit peel and aril, the edible part of the pomegranate fruit[10], and ovule developmental biology (supplemental note).

Genomic resources, which have great potential value for both basic research and crop improvement, are currently very limited for pomegranate. We have therefore sequenced and assembled the genome of *P. granatum* ‘Taishanhong’, a widely grown cultivar in China that has a bright red-colored fruit at the ripe stage. This genome sequence represents the first reference in the Lythraceae family. Genome and transcriptome analyses presented in this study provide insights into the pomegranate taxonomic status and evolution, as well as the molecular mechanisms underlying ellagitannin-based compound metabolism, anthocyanin biosynthesis, and ovule development.

## Results

### Genome sequencing and assembly

We used the whole-genome shotgun sequencing approach to generate ~67 Gb of high-quality sequences (supplemental table S1), representing approximately 200 X coverage of the pomegranate genome, which has an estimated size of 336 Mb, based on the K-mer depth distribution analysis of the sequenced reads (supplemental fig. S1) and the flow cytometry analysis (supplemental table S2). The final assembled sequence was 274 Mb, representing 81.5% of the pomegranate genome. The assembly consisted of 2,177 scaffolds (≥ 1kb) with an N50 of 1.7 Mb and 7,088 contigs with an N50 of 97 kb (table 1, supplemental table S3). The GC content of the assembled pomegranate genome was 39.2%, similar to that of *Eucalyptus grandis,* which is the evolutionarily closest species of pomegranate that has genome sequenced[11].

**Table 1.** Statistics of pomegranate genome assembly and annotation

|  |  |
| --- | --- |
| Estimated genome size (Mb) | 336 |
| Number of scaffolds ($\geq 100$ bp) | 2,117 |
| Total size of assembled scaffolds (Mb) | 274 |
| N50 scaffold length (Mb) | 1.7 |
| Longest scaffold (Mb) | 7.6 |
| Number of contigs ($\geq 100$ bp) | 7,088 |
| Total size of assembled contigs (Mb) | 269 |
| N50 contig length (Kb) | 97.0 |
| Largest contig (Kb) | 528.6 |
| GC content (%) | 39.2 |
| Number of gene models | 30,903 |
| Mean transcript length (bp) | 2,332.8 |
| Mean coding sequence length (bp) | 1,110.4 |
| Mean number of exons per gene | 4.52 |
| Mean exon length (bp) | 245.9 |
| Mean intron length (bp) | 347.6 |

A quality evaluation using BUSCO[12] revealed that 97.7% of the core eukaryotic genes were captured by the pomegranate genome assembly and that 96.2% were complete. In addition, our assembled sequence covered >99% of the 2,397 pomegranate expressed sequence tags (ESTs) downloaded from GenBank (supplemental table S4). Finally, our assembled genome covered >94% of the unigenes assembled from our pomegranate RNA-Seq data (supplemental table S5). Taken together, these results indicate that the assembled pomegranate genome sequence is of high quality.

### Genome annotation

Repetitive sequences generally constitute a large portion of a plant genome and can contribute to the plant genome evolution due to the roles in both genome size variation and functional adaption[13]. The repetitive DNA accounted for 51.2% (140.2 Mb) of the genome assembly (Table 1), a greater amount of the genome than in similarly sized plant genomes, including *Fragaria* vesca[14]. Indeed, 82.1% of pomegranate repetitive sequences could be annotated as transposable elements (TEs), of which the long terminal repeat (LTR) elements composed the most abundant (Supplemental Table S6). Among five sequenced resides species (supplemental fig. S3), the fraction of the genome contributed by LTR retrotransposons increases with genome size from *Arabidopsis thaliana* [15], the smallest genome (~15% of its 125-Mb genome consists of LTR retrotransposons), to pomegranate (17.4% retrotransposons) and *E. grandis(~640* Mb, ~20.7% retrotransposons). Clearly, the LTR dynamics is a major contributor to the 1C value differences among plants [13–16]. The two major subfamilies of LTR families found in pomegranate genome are Copia (~5.87% of total TEs) and Gypsy (~11.55%) (supplemental table S6), which differ in the order of pol encodes protease (PR), reverse transcriptase (RT), ribonuclease H (RH), and integrase (INT) domains in the polyprotein (Gypsy: PR-RT-RH-INT, Copia: PR-INT-RT-RH)[16]. Kimura distances (K-values; Kimura 1980) for all Copia and Gypsy copies were characterized to estimate the “age” and transposition history of Copia and Gypsy (supplemental fig. S4). Pomegranate genome only underwent a more recent expansion of Copia and Gypsy. Conversely, ancient divergent Copia and Gypsy elements with high K-values as well as recent activity with low K-values were found in *V. vinifera,* a reference genome of eudicots ancients. Kimura profiles consistently supported that Copia and Gypsy retrotransposons existed early in the angiosperm history and diverged into heterogeneous subgroups before the modern plant orders arose [13]. Moreover, the divergent fraction of LTR members is responsible for special biology processes, such as the regulation of pigments biosynthesis and ovule development in plants[17]. Copia-induced alleles of several genes such as *flavonoid 3’-hydroxylase (F3’H)* and *dihydroflavonol 4-reductase (DFR)* alter the expression patterns in the anthocyanin biosynthesis. Transposable elements in promoters of *AINTEGUMENTA* (ANT)[18] were inferred to be related with ovule development. Using RNA-Seq, we found that TPMs of Copia, Gypsy and LARD copies were significantly (p<0.001) increased as the development of peels or arils, which might be responsible for the anthocyanin biosynthesis (supplemental fig. S5).

We predicted a total of 30,903 protein-coding genes in the pomegranate genome, with a mean coding sequence length of 1,110 bp and 4.5 exons per gene (Table 1). Of these genes, 89% could be annotated using the GO[19], KEGG[20], TrEMBL[21], COG[22], or the GenBank nr databases (supplemental table S8). Furthermore, conserved domains in 80% of the predicted protein sequences were identified by comparing them against the InterPro database[23]. In addition to the protein-coding genes, 601 miRNA, 54 rRNA and 144 tRNA genes were also identified in the pomegranate genome (supplemental table S9).

### Comparative genomic analysis between pomegranate and other plant species

A gene family cluster analysis of the complete gene sets of pomegranate, *E. grandis*, apple (*M. domestica),* Arabidopsis *(Arabidopsis thaliana)* and grape was performed. A total of 22,426 genes in the pomegranate genome were grouped into 13,747 gene clusters, of which 8,459 were shared between all the five species (fig. 1A). Pomegranate shared more gene family clusters with *E. grandis* (11,992) than with any of the other three species, and we also inferred a relatively close taxonomic relationship between these two species from their presence in a shared clade in a phylogenetic tree constructed with 172 single-copy genes (supplemental fig. S2). Furthermore, we assembled the transcriptomes of six species in the Lythraceae family, as well as that of *Oenothera biennis* (Onagraceae family of the order Myrtales), and then reconstructed a species tree of the Lythraceae family (fig. 1B). On the basis of this tree, four pomegranate cultivars and *Lagerstroemia indica* were classified into one monophyletic clade, and then clustered in a group with two species from the *Cuphea* genus. Based on the genomic phylogenetic analysis, we concluded that the *Punica* genus belongs to the Lythraceae family.

**Figure 1.**
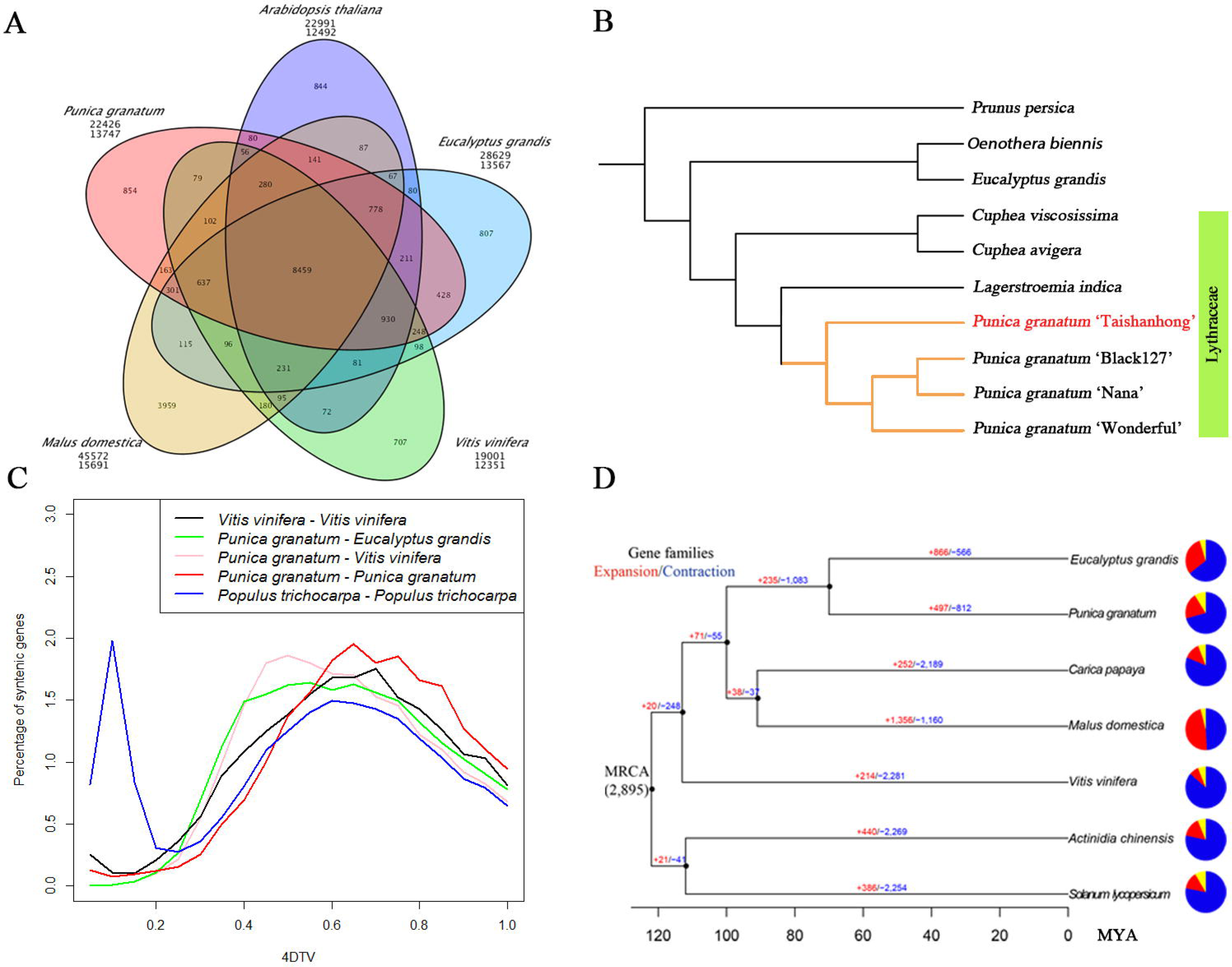
Comparative genomic analysis of pomegranate and other eudicot species. (**A**) Venn diagram of shared orthologous gene families in pomegranate, *Eucalyptus grandis, Malus domestica, Vitis vinifera,* and *Arabidopsis thaliana.* The gene family number is listed in each component. (**B**) Phylogenetic tree constructed from 106 single-copy gene families. (**C**) Distribution of the 4DTv distance between syntenically orthologous genes. (**D**) Gene family expansion and contraction analysis. MRCA, most recent common ancestor. Gene family expansions and contractions are indicated by numbers in red and blue, respectively. Blue and red portions of the pie charts represent the contracted and expanded gene families relative to MRCA, respectively, while the grey portions represent the conserved gene families.

We identified 2,749 syntenic blocks within the pomegranate genome, and also identified syntenic blocks between the genomes of pomegranate and grape, and pomegranate and *E. grandis,* as well as within the grape and *Populus trichocarpa* genomes. The distribution of 4DTv (transversions at fourfold degenerate sites) of homologous gene pairs within these syntenic blocks suggested that pomegranate has not undergone any recent lineage-specific whole genome duplication (WGD) events, but shared the paleohexaploidy event (γ) of all eudicots (fig. 1C). However, the divergence between pomegranate and *E. grandis,* estimated based on the MCMCtree[24], occurred at ~69.6 (51.5-85.0) million years ago (MYA), which is after the paleotetraploidy event (109.9 MYA) identified in the *E. grandis* genome[11] (fig. 1D), indicating that this WGD event is shared by pomegranate and *E. grandis.* Further analysis of the syntenic blocks between pomegranate and grape, whose genome has not undergone recent genome duplication[25], and pomegranate and *E. grandis* suggested that the majority of grape syntenic regions had two orthologous regions in pomegranate, while the majority of *E. grandis* syntenic regions had one in pomegranate (supplemental table S9). In addition, Ks (number of synonymous substitutions per synonymous site) values of paralogous genes from the ancient duplications within pomegranate and *E. grandis* showed similar distribution patterns (supplemental fig. S6). Taken together, these findings strongly support that the paleotetraploidy event identified in *E. grandis* is shared by pomegranate.

We identified 15 gene families that have undergone significant (p-value < 0.01) expansion in the pomegranate genome. These families were found to be enriched with genes involved in self-incompatibility and other specialized biological pathways (supplemental fig. S7), suggesting that these pathways have evolved distinctly in pomegranate compared to other plant species.

### Biosynthesis of unique ellagitannin-based compounds

Pomegranate fruit have much higher antioxidant activity than apple, orange, grape and other fruits[2], which is attributed to their high content of polyphenols; specifically punicalagins, punicalins, gallagic acid, ellagic acid, and other ellagitannin-based compounds. Notably, the results of human clinical trials suggest that these compounds can contribute to reduced rates of cardiovascular disease, diabetes, and prostate cancer[3]. To investigate the molecular mechanisms underlying the biosynthesis of the ellagitannin-based compounds, we performed integrated genomic and transcriptomic analyses of genes in the ellagitannin biosynthetic pathway (fig. 2A, supplemental note).

**Figure 2.**
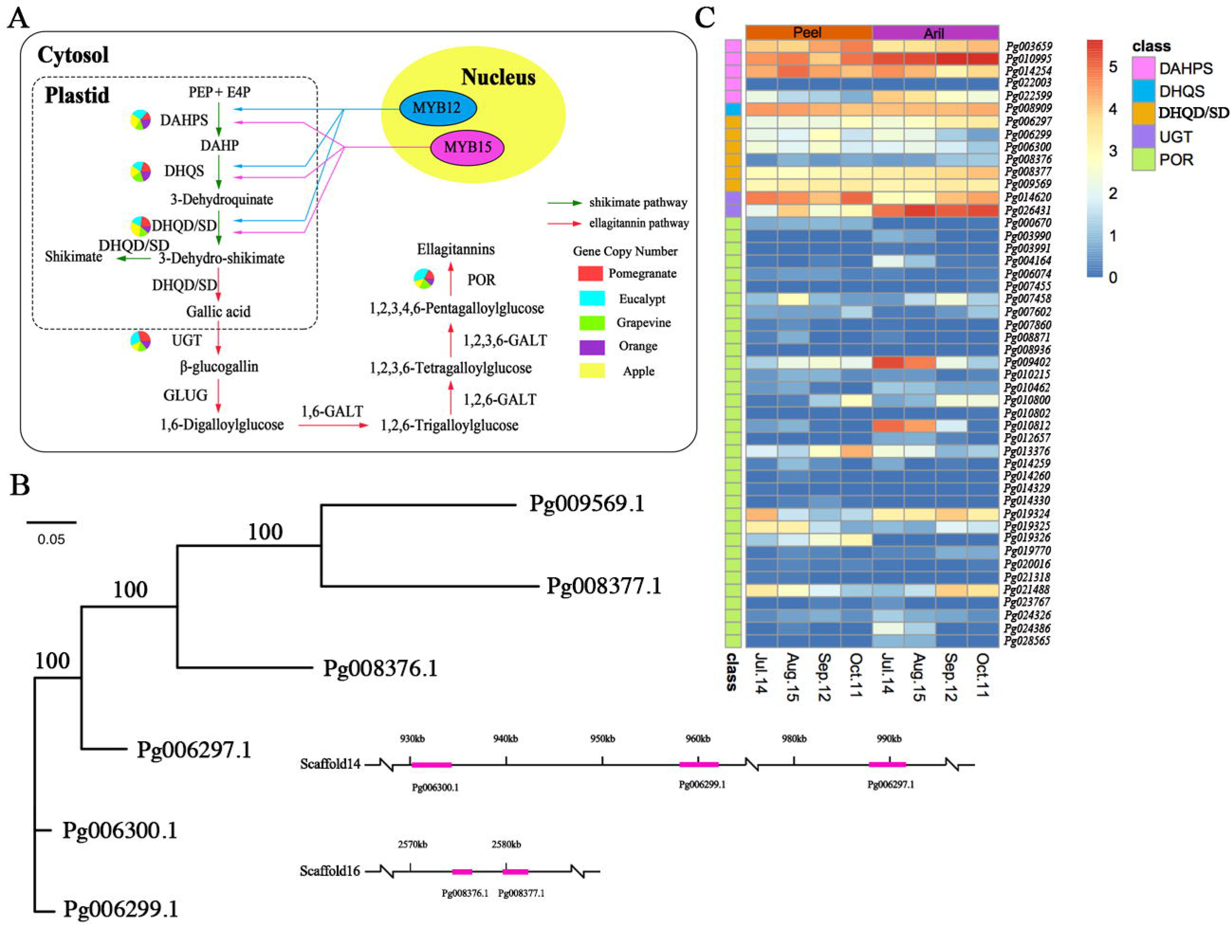
Evolution of ellagitannin biosynthesis in pomegranate. (**A**) The ellagitannin biosynthetic pathway. Green and red arrows represent the shikimate and ellagitannin pathways, respectively. The numbers of genes in each family in the ellagitannin metabolic pathway in pomegranate, *Eucalyptus grandis*, grape, orange, and apple genomes are shown in the pie charts. (**B**) Phylogenetic analysis and genome locations of DHQD/SD genes in pomegranate. (**C**) Expression heat map of genes related to the synthesis of ellagitannins in peel and aril during pomegranate fruit development.

The enzyme 3-dehydroquinate dehydratase/shikimate dehydrogenase (DHQD/SD) serves as a key bridge linking the shikimate pathway and ellagitannin biosynthetic pathway[26]. Six *DHQD/SD* genes were identified in the pomegranate genome, of which three *(Pg006297.1, Pg006299.1* and *Pg006300.1)* were tandemly duplicated and located in a 100-kb region (fig. 2B). Although all three of these genes were highly expressed in both fruit peels and arils, *Pg006299.1* and *Pg006300.1* showed decreasing expression during fruit development (fig. 2C, supplemental fig. S5), consistent with the fact that levels of pulicalagin, ellagic acid and gallic acid also decrease during pomegranate fruit development[27], indicating their potential important roles in ellagitannin biosynthesis. Two other *DHQD/SD* genes, *Pg008377.1* and *Pg008376.1,* were also tandemly duplicated. *Pg008377.1* was highly expressed in fruits while *Pg008376.1* had a very low expression level (fig. 2C), suggesting their subfunctionalization after tandem duplication. In addition, two *UDP-glucose:gallate glucosyltransferase (UGT)* genes *(Pg014620.1* and *Pg026431.1)* were identified in the pomegranate genome and they showed distinct expression patterns: *Pg014620.1* was expressed higher in peel than in aril, while *Pg026431.1* was expressed higher in aril than in peel (fig. 2C; supplemental fig. S8), suggesting the tissue-specific roles of these two genes in ellagitannin biosynthesis.

Another key enzyme family in the ellagitannin biosynthetic pathway is pentagalloylglucose oxygen oxidoreductase (POR). A total of 34 *POR* genes were identified in the pomegranate genome (supplemental table S11). Phylogenetic analysis placed these genes into twelve groups and member expansion was observed in group 1 (supplemental fig. S9). Four genes in group 1 *(Pg007458.1, Pg019324.1,*

*Pg019325.1* and *Pg021488.1)* were highly expressed in both fruits and arils, and the expression of *Pg019324.1* and *Pg019325.1* showed a decreased trend in the peel during fruit development (supplemental fig. S8). Of the genes in the other groups, *Pg009402.1* and *Pg010812.1* showed a clear descending expression trend in aril during fruit development (supplemental fig. S8). Reduced expression of these genes during peel and aril development could be responsible for the decreased productions of pulicalagin, ellagic acid and gallic acid[28].

Interestingly, sequence homology searches did not reveal any genes predicted to encode βxs-glucogallin O-galloyltransferase (GLUG) or galloyltransferase (GALT), which are also enzymes in the ellagitannin biosynthetic pathway. These genes may have diverged to a degree in pomegranate that sequence homology has been lost or pomegranate may have developed alternative reactions for the steps catalyzed by these two enzymes.

### Evolution of the anthocyanin biosynthetic pathway

Anthocyanins are the major pigments and responsible for the color of pomegranate fruits[29]. Differed from other fruits such as *Litchi chinensis[30]* and *V.* vinifera[31], both peel and aril are bright red at the ripe stage in pomegranate (fig. 3A). Although the anthocyanin biosynthetic pathway in fruit peels has been studied in several fruit tree species[32], it has not been well characterized in arils. From our genome assembly, 20 anthocyanin biosynthesis genes from 9 gene families were identified, including 11 for anthocyanidin synthase, and 9 for glycosylation and methylation (fig. 3B, supplemental table S12). The wider diversity of anthocyanin compounds comes from the glycosylation[33] and methylation[34] of the basic flavonol structure. In anthocyanidin synthetic pathway, each enzyme had substantial expression in both peel and aril, and the most members had preferential expression in peel (fig. 3C and D, supplemental fig. S10). By contrast, for anthocyanidin modification only three genes (Pg010555.1, Pg002351.1, Pg021629.1) highly expressed in peel (fig. 3C and D). High performance liquid chromatography analyses show that the total anthocyanin contents in peel (~118.65 mg/100g) are higher than that in aril (~36.41 mg/100g)[27]. Our results support the tissue-specific expression pattern for anthocyanin biosynthesis. Integrated RNA-Seq and iTraq analyses indicate that highly up-regulated expression of enzymes such as chalcone synthase (CHS), chalcone isomerase (CHI), flavonoid 3-hydroxylase (F3H), flavonoid 3’-hydroxylase (F3’H), dihydroflavonol 4-reductase (DFR), anthocyanidin synthase/leucoanthocyanidin dioxygenase (ANS/LDOX), UDP-glucose:flavonoid glucosyltransferases (UFGT), and anthocyanin O-methyltransferase (AOMT) are responsible for the skin gradually changing from white to red (fig. 3A)[35].

**Figure 3.**
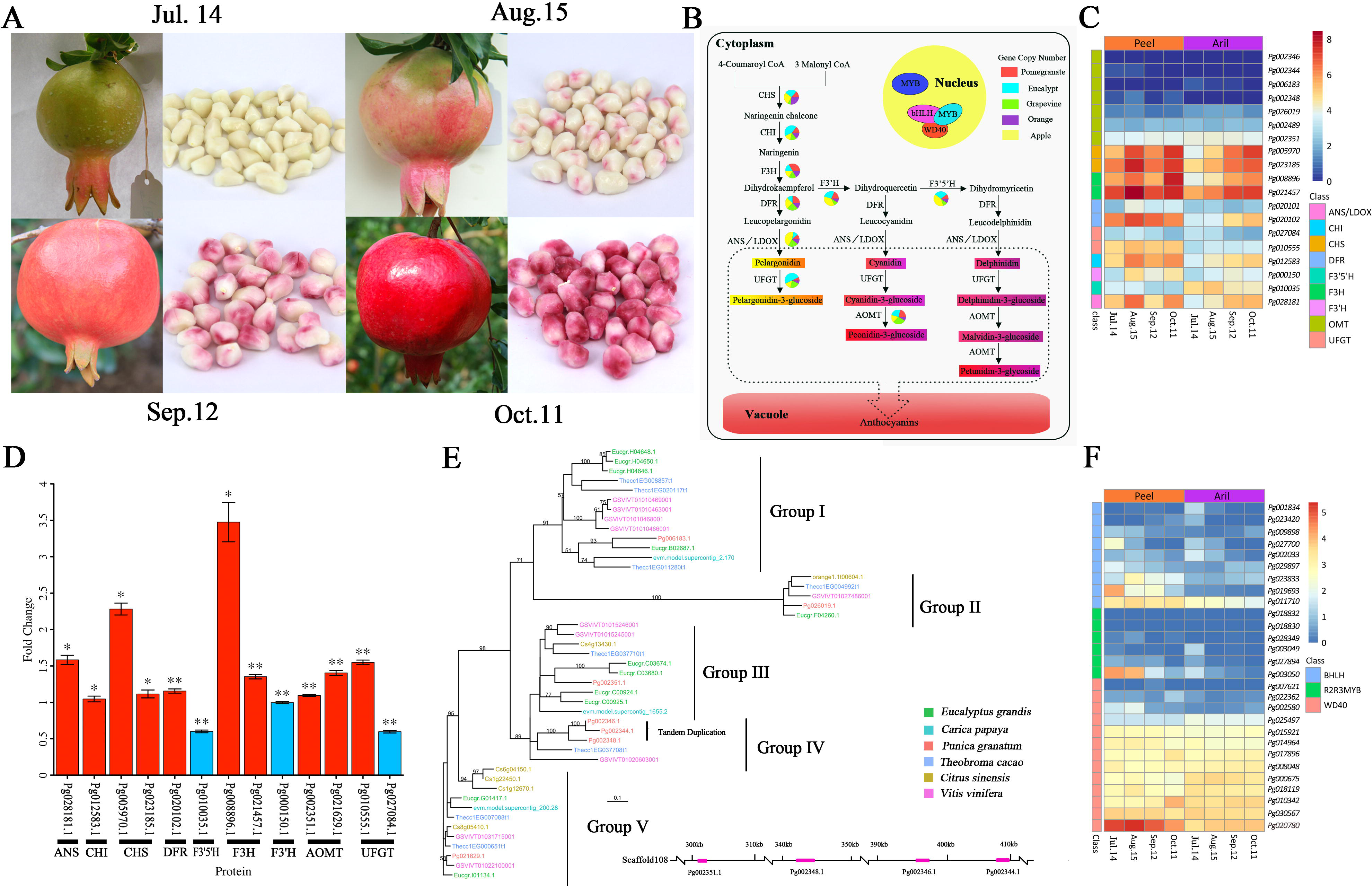
Anthocyanin biosynthetic pathway in pomegranate. (**A**) Fruits and arils of ‘Taishanhong’ pomegranate at different developmental stages. (**B**) The anthocyanin biosynthetic pathway. The numbers of genes in each family in the anthocyanin biosynthetic pathway in pomegranate, *E. grandis*, grape, orange and apple are shown in the pie charts. (**C**) Expression heat map of genes related to the synthesis of anthocyanins in peel and aril during fruit color development. (**D**) Expression change of proteins from Jul.14 to Oct.11. The red and blue bars represent up-and down-regulated proteins, respectively. Values present means±s.e. of three repeats. **P<0.01, *P<0.05 by Student’s t-test. (**E**) Phylogenetic analysis of the AOMT genes in plants within the Malvids clade and the outgroup species, grape. Top: four putative tandemly duplicated pomegranate AOMT genes located in scaffold 108 are shown as purple bars. (**F**) Expression heat map of the putative MBW complex genes.

The pomegranate and grape genome have the same copy (7) of anthocyanin O-methyltransferase (AOMT) genes which are higher than other three species (Supplemental table S12). Different copy numbers indicate an obvious expansion of this enzyme family in pomegranate. The gene number of the AOMT family does not seem to correlate with genome size and chromosome number of a given species. *E. grandis* (640 Mb, 2n=22)[11] only has 6 genes, and *C. sinensis* (367 Mb, 2n=18)[36] only has 2 genes. AOMTs play the final role in anthocyanin biosynthesis pathway, mediating the methylation of anthocyanins[34]. High copy number of AOMT genes in fruit species with diverse anthocyanins supports a putative link between expansion of the AOMT family and the ability to produce anthocyanin.

Phylotranscriptomic analysis shows that AOMT enzymes obviously expanded in ‘Taishanhong’ than other two pomegranate cultivars ‘Nana’ and ‘Black127’ (supplemental fig. S11). The divergent AOMTs are inferred to be responsible for distinct colors of pomegranate fruits. Phylogenetic analysis of pomegranate AOMTs and their homologs from six other plant species within the Malvidae clade revealed one recent *AOMT* gene expansion in the pomegranate genome, comprised of three tandemly duplicated genes (fig. 3E). These three genes were under relaxed purifying selection, as indicated by their low dN/dS values (supplemental table S13), and they showed low expression levels in peel and aril (supplemental fig. S10). These seven AOMT enzymes exhibit distinct expression patterns during fruit development. RNA-Seq and iTraq analyses indicate that of the seven AOMT gene expressions, only *Pg002351.1* and *Pg021629.1* were highly upregulated in peel (supplemental fig. S10, fig. 3D). *Pg002351.1* upregulated and *Pg021629.1* downregulated during fruit development in aril. The tissue-specific expression pattern of AOMTs are responsible for anthocyanin in peel and aril. Pg002351.1 and Pg021629.1 are classed into Groups III and V, respectively (fig. 3E). Integrated phylogenetic, transcriptomic and proteomic analyses conclude that AOMT Group III might have evolved independently towards functioning in anthocyanin biosynthesis.

F3’H and F3’5’H are responsible for the production of cyanidin-and delphindin-based anthocyanins, respectively[37–38]. The expression of the *F3’H* gene *(Pg000150.1)* was higher than that of the *F3’5’H* gene *(Pg010035.1)* (Supplemental Fig. S10), indicating that di-hydroxylated anthocyanins are the main pigments in pomegranate fruits, as previously reported [35]. The cyanidin-and delphindin-based anthocyanins can be further modified by AOMT enzymes (Fig. 3B). *Pg002351.1 (AOMT)* and *Pg000150.1 (F3’H)* showed similar expression patterns during fruit development, while *Pg021629.1 (AOMT)* and *Pg010035.1 (F3’5’H)* showed similar trends (Supplemental fig. S10), implying that *Pg002351.1* and *Pg021629.1* might be responsible for the methylation of cyanidin-and delphindin-based anthocyanins, respectively.

Anthocyanin biosynthetic genes are activated by a transcriptional activation complex (the MBW complex) consisting of R2R3-MYB, BHLH, and WD40 proteins[32]. In *Arabidopsis,* genes encoding enzymes in the early steps of the anthocyanin biosynthetic pathway that lead to the production of flavonols, are activated by three R2R3-MYB regulatory genes *(AtMYB11, AtMYB12* and *AtMYB111),* whereas the activation of the late biosynthetic genes, leading to the production of anthocyanins, requires an MBW complex[39]. In the pomegranate genome, we identified six *R2R3-MYB* genes, nine *BHLH* genes, and thirteen *WD40* genes that were highly expressed in the peel and aril (Fig. 3F), suggesting their roles in regulating anthocyanin production in pomegranate fruit. Recently, an identified NAC transcriptional factor BLOOD (BL) up-regulates the accumulation of anthocyanin in peach[40]. However, blast searching yielded no significant hits in our genome assembly, indicating a discrepancy in the mechanisms of anthocyanin biosynthesis between these two species.

### Ovule developmental biology

The polycaryoptic trait is a common target of the plant breeding programs. In pomegranate, more than one hundred ovules can grow in a single ovary, and they develop into seeds with arils, which consist of epidermal cells derived from the integument[41]. Compared to cucumber and tomato, which have parietal[42] and axial[43] placentas, respectively, pomegranate carpels become superposed into two or three layers by differential growth, the lower comprised of axial placentas and the upper ostensibly parietal placentas[4]. Consequently, pomegranate represents a unique system for studying ovule developmental biology.

We identified and compared genes involved in ovule development from the genomes of pomegranate, castor bean (another species with arils[44]), cucumber and tomato (Fig. 4A). The pomegranate genome has 237 candidate genes belonging to the twelve families affiliated with ovule development (Fig. 4B). The AG-clade, including the AG, SEP, SHP, and STK families, had the largest copy number (39) in the pomegranate genome (Fig. 4B). AG-clade genes are required for specifying the ovule identity[18–45], suggesting that the expansion of AG-clade genes might play an important role in the development of the pomegranate-specific type of ovules. Furthermore, structure, transcriptome, and proteome analyses showed that the BEL1 gene (Pg029909.1) occurred frame-shift and functionally inactivated, which had low transcriptional expression and none of peptide expression. BEL1 gene had negative role in regulating WUS expression, resulting in carpelloid structures[18]. Low copy of BEL1 genes and pseudegenization suggest a possible links between contraction and inactivation of BEL1 family and multi-carpels formation. Additionally, the pomegranate genome also has a higher copy number (87) of *CUC* genes than the other three genomes (Fig. 4B). CUC proteins have reported to regulate ovule production[46], and expansion of the CUC family in the pomegranate genome may be a key factor in the production of the large number of ovules[46]. Based on our comparative genomic analysis, the pomegranate-specific ovule development and the polycaryoptic phenotype can likely be attributed to the expansions of the AG and CUC families and contraction and inactivation of BEL1 family.

**Figure 4.**
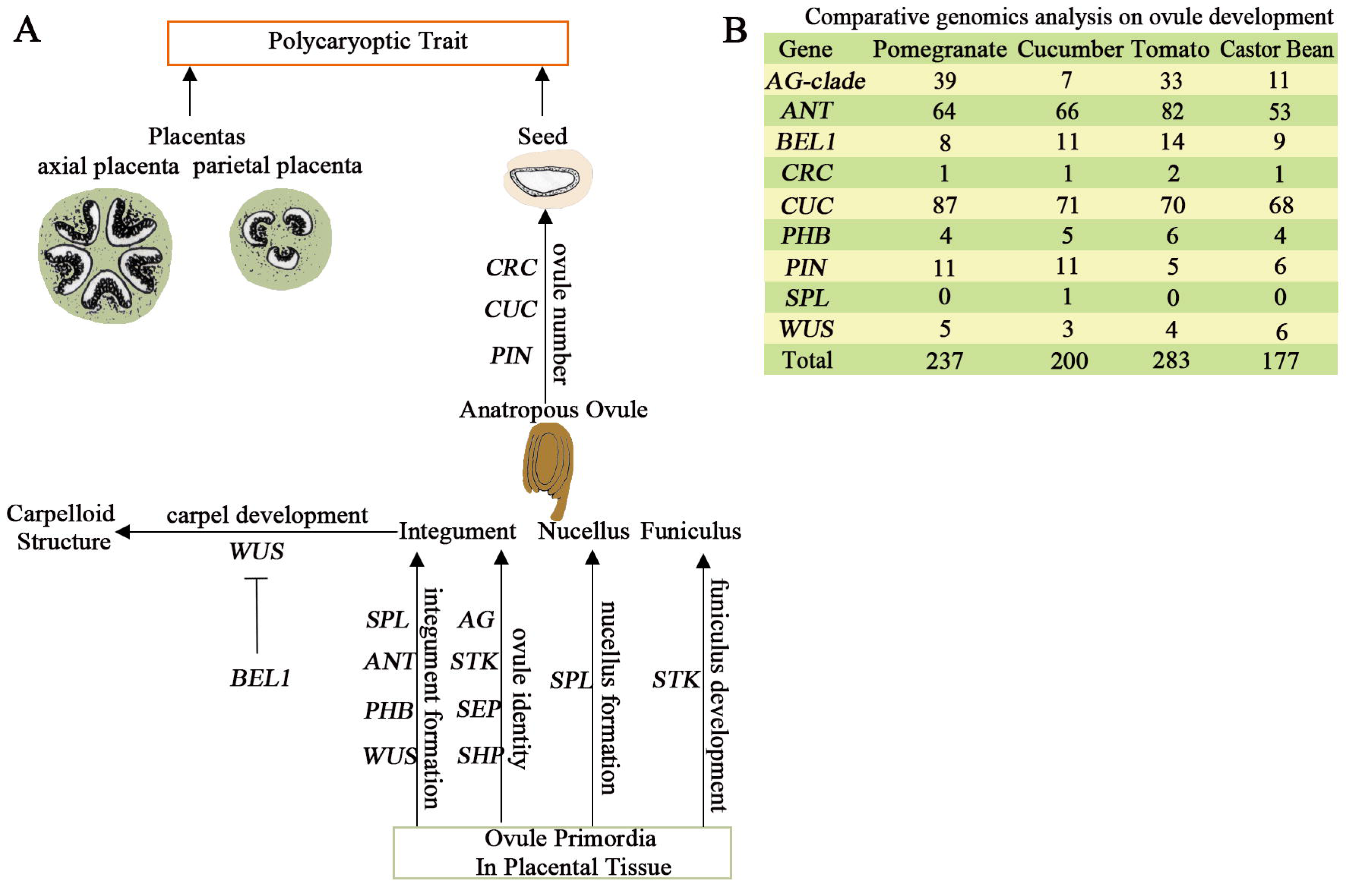
Regulation of ovule development in pomegranate. (A) Ovule development. Genes involved in ovule development include those belonging to MADS-box gene families, such as AGAMOUS (AG), SEEDSTICK (STK), SEPALLATA (SEP) and SHATTERPROOF (SHP); HOMEOBOX gene families, such as WUSCHEL (WUS), PHABULOSA (PHB) and BELL1 (BEL1); AP2-like gene families, such as AINTEGUMENTA (ANT); NOZZLE gene families, such as SPOROCYTELESS (SPL); YABBY gene families, such as CRABS CLAW (CRC); and other families, such as CUP-SHAPED COTYLEDONS (CUC) and PIN-FORMED (PIN). (B) Comparative analysis of gene families involved in ovule development. The AG-clade contains AG, STK, SEP, and SHP genes.

## Discussion

A high-quality reference genome sequence of pomegranate was assembled, which offers a valuable resource for resolving the previously debated taxonomic status of the *Punica* genus [7]. *Punica* was previously considered a member of the monogeneric Punicaceae family[47] but was later moved into the Lythraceae family[7–8]. Our phylogenomic analysis strongly supports this reclassification of *Punica* into Lythraceae. Consequently, pomegranate represents the first species in the Lythraceae family that has a sequenced genome, providing an important reference for future comparative genomics and evolutionary studies.

Pomegranate fruits are highly enriched in ellagitannin-based compounds, which are known to possess antioxidant activities[3]. Another important fruit quality trait of pomegranate is the color formation related to the anthocyanin biosynthesis in the peel and aril. Our genomic, RNA-Seq and iTraq analyses of the pomegranate offer deeper insights into the molecular mechanisms underlying ellagitannin and anthocyanin biosynthesis. Several key gene families in each of the associated pathways were found to have undergone tandem duplications and specific family members showed differential expression patterns in the peel and/or aril during fruit development, indicative of their important roles in the production of these compounds.

With hundreds of ovules in a single ovary[4], rare heterotypic placentation, and arils developed from integuments[41], pomegranate has provided a unique system for studying the ovule development. Our comparative genomic analysis provided evidence that the pomegranate-specific ovule development and the polycaryoptic phenotype can be attributed, at least in part, to the expansions of the MADS-box AG clade and the CUC family, respectively.

In summary, pomegranate genomic information represents an invaluable resource for the genetic improvement of the crop and for better understanding of genome evolution. Genetic markers can be developed based on this genome sequence for studies involving genetic map construction, positional cloning, strain identification and marker-assisted selection, which will collectively accelerate pomegranate breeding.

## Materials and Methods

### Sample preparation and sequencing

Genomic DNA was extracted from the leaves of *P. granatum* ‘Taishanhong’, using the CTAB protocol. Paired-end and mate-pair Illumina genome libraries with insert sizes ranging from 220 bp to 17 kb were constructed using the NEB Next Ultra DNA Library Prep Kit (NEB, USA) and sequenced on a HiSeq 2500 system (Illumina, USA) according to the manufacturer’s instructions. Raw reads were processed to remove low quality and adaptor sequences, and to collapse duplicated reads using NxTrim[48].

### De novo genome assembly

The high-quality cleaned reads were assembled *de novo* using ALLPATHS-LG[49], and the mate-pair reads were then used to construct scaffolds, using SSPACE2.0 [50]. Gap filling was performed using GapCloser provided in SOAPdenovo2 [51]. Assembled scaffolds were compared against the Genbank nt database using megablast and against a set of known microbial proteins using BLASTX. Scaffolds classified as microbial sequences, unanchored rDNA, mitochondrion, chloroplast, and repetitive sequences, as well as those < 1 kb were removed from the final assembly.

### Repeat annotation

We first identified repeat sequences in the *P. granatum* genome using the *de novo* prediction programs, LTR_FINDER[52], MITE-Hunter[53], RepeatScout[54] and PILER-DF[55], and then classified the identified repeat sequences with PASTEClassifier (v1.0)[56]. The classified repeat sequences and the Repbase database[57] were combined to construct a non-redundant repeat sequence library. RepeatMasker (v4.0.6; http://www.repeatmasker.org) was used to identify the *P. granatum* repeat sequences based on the constructed repeat sequence library.

### Gene prediction and annotation

The repeat-masked *P. granatum* genome sequence was used for gene prediction with the following methods: (i) *de novo* gene prediction, (ii) homologous sequence searching, (iii) transcriptome sequence mapping. We first assembled the RNA-Seq reads into contigs using Trinity (v2.1.1)[58]. The *P. granatum* specific parameter file was trained by the *de novo* gene prediction software Augustus (v1.0.2)[59] using the *bona fide* gene models, which were identified from the assembled RNA-Seq contigs by PASA (v1.2)[60]. Using this parameter file, we performed *de novo* gene predictions using Augustus, SNAP[61] and GlimmerHMM (v0.5.9)[62], respectively. We also performed *de novo* gene predictions using Genscan (v0.5.9)[63] and GeneID (v1.4)[64] with the Arabidopsis parameter file. In homologous sequence searches, we aligned the protein sequences from *E. grandis,* the plant specific UniProtKB/Swiss-Prot database[65] and the GenBank nr database against the *P. granatum* genome using TBLASTN with a sequence identity >50% and an E-value cutoff of 1E-5. GeneWise[66] was then used to extract the accurate exon-intron information. GMAP (v1.0.0)[67] was used to align the assembled RNA-Seq contigs to the *P. granatum* genome. Finally, we generated an integrated gene set using GLEAN[68].

Functional annotation of the predicted genes was performed by comparing their protein sequences against a number of protein sequence databases, including GenBank nr, COG[22], KEGG[20] and TrEMBL[21], using BLASTP with an E-value cutoff of 1E-5.

### Collinearity and WGD

All-against-all BLASTP analyses of protein sequences were performed between *P. granatum, V. vinifera, E. grandis* and *P. trichocarpa* using an E-value cutoff of 1E-10. Syntenic regions within and between species were identified using MCScan[69], based on the BLASTP results. A syntenic region was identified if it contained a minimum of 10 and a maximum of 25 genes in the identified gene pairs. Protein sequences of homologous gene pairs in the identified syntenic regions were aligned by MUSCLE (v3.8.31)[70], and the protein alignments were then converted to coding sequence (CDS) alignments. The Ks value of each gene pair was calculated using the Yn00 program in the PAML package[24] and the 4DTV value of each gene pair was calculated using the sum of transversions of four-fold degenerate sites divided by the sum of fourfold degenerate sites and then corrected using the HKY model[71].

### Gene family evolution and phylogenetic analyses

Protein sequences of *P. granatum, E. grandis, M. domestica, A. thaliana* and *V. vinifera* were used in an all-against-all BLASTP analysis. The results were analyzed using the OrthoMCL software[72] with an MCL inflation parameter of 1.5 to identify gene family clusters. Gene family clusters were also identified among *P. granatum, E. grandis, M. domestica, C. papaya, V. vinifera, S. lycopersicum* and *A. chinensis.* Single copy gene clusters shared by all 7 species were identified and used to construct a phylogenetic species tree using PhyML (v3.0)[73]. The divergence time was estimated by MCMCtree[24] using the known divergence time of *V. vinifera* and *M. domestica,* and *V. vinifera* and *A. chinensis* from the TimeTree database[74]. In addition, we also used a Pfam domain-based method to infer the gene family expansions as described in Albertin et al (2015)[75].

To determine the taxonomic position of *P. granatum,* transcript assemblies were performed using Trinity[58] and RNA-Seq reads from other *P. granatum* cultivars and other species (three cultivars of *P. granatum:* ‘Black127’, ‘Nana’, and ‘Wonderful’; three species from the Lythraceae family: *Lagerstroemia indica, Cuphea viscosissima,* and *Cuphea avigera* var. *pulcherrima*; and one species from the Onagraceae family: *Oenothera biennis)* downloaded from the NCBI sequence archive (SRA) database (Accession numbers: SRX395468, SRX395465, SRX034876, SRX470007, SRX1361461, SRX1361546, ERX651036, ERX651029, ERX651035, ERX651028 and ERX651064). The open reading frame (ORF) of each unigene was identified using GeneMarkS-T[76] and the translated amino acid sequences were then used for phylogeny reconstruction.

### RNA collection and sequencing

Peel and aril samples were collected from four different developmental stages (on Jul. 14, Aug. 15, Sept. 12 and Oct. 11, respectively) of pomegranate fruits. Three biological replicates were analyzed for each sample. Total RNA was extracted using TRI Reagent (Sigma Life Science, USA) according to the manufacturer’s instructions. RNA-Seq libraries were constructed using the NEB Next UltraTM RNA Library Prep Kit (NEB, USA) and sequenced on an Illumina HiSeq 4000 platform (Illumina, USA) according to the manufacture’s protocols.

### Quantification and differential gene expression analysis

Paired-end RNA-Seq reads were processed to remove adaptor sequences and low-quality reads and then mapped to the *de novo* assembled pomegranate genome sequence using TopHat2 (v2.0.13)[77] with default parameters. Cufflinks (v2.2.1)[77] was used to assemble the mapped reads for each sample. The assembled contigs were then merged with the reference gene annotation into a unified annotation, which was used to quantify gene expression in each sample. We used the FPKM (fragments per kilobase exon model per million mapped fragments) as the normalized expression level of genes. Differentially expressed genes between different tissues or across different developmental stages were identified using DESeq (v1.20.0)[78]. Genes with an adjusted P-values <0.01 were considered to be differentially expressed.

### iTraq assays

Peels from Jul. 14 and Oct. 11 were ground in liquid nitrogen. Proteins were precipitated according to the reference methods[79]. Desalted peptides were then labeled with iTRAQ reagents (Applied Biosysterms) according to the manufacturer’s instructions. Nano-HPLC-MS/MS analysis was performed on a nanoAcquity system (Waters) connected to an LTQ-Orbitrap XL hybrid mass spectrometer (Thermo Electron) equipped with a PicoView nanospray interface (New Objective). iTRAQ 8plex was chosen for quantification during the search simultaneously.

### Accession codes

The pomegranate whole-genome sequence has been deposited in GenBank under a BioProject with accession number PRJNA355913. The data will be made public once the manuscript is accepted.

## Acknowledgements

This work was supported by the National Natural Science Foundation of China (31272143), the Initiative Project for Talents of Nanjing Forestry University (GXL2014070), the Priority Academic Program Development of Jiangsu High Education Institutions (PAPD), the Research Fund for Postgraduate Innovation Project of Jiangsu Province (KYLX16_0857), the Doctorate Fellowship Foundation of Nanjing Forestry University, and the United States National Science Foundation (IOS-1539831).

### Author contributions

Z.Y., Y.F., T.Z., Z.F., and F.H. contributed equally to this work. T.Z., C.L., W.X., H. X., J.Z., Y.L., Y.C., M.Y., X.H., and H.W. contributed to plant sample collection; T.Z., F.H., M.L., W.Z., D.G., Y.X., and D.L. worked on genomic DNA sequencing and genome assembly; T.Z., Z.F., F.H., C.L., W.X., W.Z., M.Z., Y.J., H.X., H.D. and L.W. contributed to transcriptome sequencing and gene expression analyses; T.Z., Z.F., F.H., W.Z., and Y.J. conducted genome annotation and comparative genomic analyses; T.Z., Z.F. and F.H. wrote and revised the manuscript; Z.Y., Y.F. and H.Z. conceived and manage the project, and designed experiments. All authors read and approved the manuscript.

## References

1 Holland D, Hatib K, Bar-Ya′akov I. Pomegranate: botany, horticulture, breeding. Hort Rev 2009;35:127–191.

2 Halvorsen BL, Holte K, Myhrstad MCW, et al. A systematic screening of total antioxidants in dietary plants. J Nutr 2002;132:461–471.

3 Johanningsmeier SD, Harris GK. Pomegranate as a functional food and nutraceutical source. Annu Rev Food Sci Technol 2011;2:181–201.

4 Teixeira da Silva JA, Rana TS, Narzary D, Verma N, Meshram DT, Ranade SA. Pomegranate biology and biotechnology: A review. Sci Hortic 2013;160:85–107.

5 Yi Z, Feng T, Zhuang H, Ye R, Li M, Liu T. Comparison of different extraction methods in the analysis of volatile compounds in pomegranate juice. Food Anal Methods 2016;9:2364–2373.

6 Graham SA, Graham A. Ovary, fruit, and seed morphology of the Lythraceae. Int J Plant Sci 2014; 175:202–240.

7 Berger BA, Kriebel R, Spalink D, Sytsma KJ. Divergence times, historical biogeography, and shifts in speciation rates of Myrtales. Mol Phylogen Evol 2016;95:116–136.

8 Byng JW, Chase MW, Christenhusz MJM, et al. An update of the angiosperm phylogeny group classification for the orders and families of flowering plants: APG IV. Bot J Linn Soc 2016;181:1–20.

9 Ono NN, Qin X, Wilson AE, Li G, Tian L. Two UGT84 family glycosyltransferases catalyze a critical reaction of hydrolyzable tannin biosynthesis in pomegranate (Punica granatum). PLoS One 2016;11:e0156319.

10 Ben-Simhon Z, Judeinstein S, Nadler-Hassar T, et al. A pomegranate (Punica granatum L.) WD40-repeat gene is a functional homologue of Arabidopsis TTG1 and is involved in the regulation of anthocyanin biosynthesis during pomegranate fruit development. Planta 2011;234:865–881.

11 Myburg AA, Grattapaglia D, Tuskan GA, et al. The genome of Eucalyptus grandis. Nature 2014;510:356–362.

12 Simao FA, Waterhouse RM, Ioannidis P, Kriventseva EV, Zdobnov EM. BUSCO: assessing genome assembly and annotation completeness with single-copy orthologs. Bioinformatics 2015;31:3210–3212.

13 Vitte C, Panaud O. LTR retrotransposons and flowering plant genome size: emergence of the increase/decrease model. Cytogenet Genome Res 2005;110:91–107.

14 Shulaev V, Sargent DJ, Crowhurst RN, et al. The genome of woodland strawberry (Fragaria vesca). Nat Genet 2011;43:109–116.

15 Initiative AG. Analysis of the genome sequence of the flowering plant Arabidopsis thaliana. Nature 2000;408:796.

16 Lee S-I, Kim N-S. Transposable elements and genome size variations in plants. Genomics & Informatics 2014;12:87–97.

17 Feschotte C, Jiang N, Wessler SR. Plant transposable elements: where genetics meets genomics. Nat Rev Genet 2002;3:329–341.

18 Colombo L, Battaglia R, Kater MM. Arabidopsis ovule development and its evolutionary conservation. Trends Plant Sci 2008;13:444–450.

19 Ashburner M, Ball CA, Blake JA, et al. Gene Ontology: tool for the unification of biology. Nat Genet 2000;25:25–29.

20 Kanehisa M, Goto S. KEGG: kyoto encyclopedia of genes and genomes. Nucleic Acids Res 2000;28:27–30.

21 Bairoch A, Apweiler R. The SWISS-PROT protein sequence data bank and its supplement TrEMBL. Nucleic Acids Res 1997;25:31–36.

22 Tatusov RL, Galperin MY, Natale DA, Koonin EV. The COG database: a tool for genome-scale analysis of protein functions and evolution. Nucleic Acids Res 2000;28:33–36.

23 Mitchell A, Chang H-Y, Daugherty L, et al. The InterPro protein families database: the classification resource after 15 years. Nucleic Acids Res 2014;43:D213–D221.

24 Yang Z. PAML 4: Phylogenetic analysis by maximum likelihood. Mol Biol Evol 2007;24:1586–1591.

25 Jaillon O, Aury J-M, Noel B, et al. The grapevine genome sequence suggests ancestral hexaploidization in major angiosperm phyla. Nature 2007;449:463–467.

26 Maeda H, Dudareva N. The shikimate pathway and aromatic amino acid biosynthesis in plants. Annu Rev Plant Biol 2012;63:73–105.

27 Zhu FZ, Yuan ZH, Zhao XQ, Yin YL, Feng LJ. Composition and contents of anthocyanins in different pomegranate cultivars. Acta Hortic 2015;1089:35–41.

28 Han LL, Yuan ZH, Feng LJ, Yin YL. Changes in the composition and contents of pomegranate polyphenols during fruit development. Acta Hortic 2015;1089:53–61.

29 Ben-Simhon Z, Judeinstein S, Trainin T, et al. A “White” Anthocyanin-less Pomegranate (Punica granatum L.) Caused by an Insertion in the Coding Region of the Leucoanthocyanidin Dioxygenase (LDOX; ANS) Gene. PLoS One 2015;10.

30 Hu B, Zhao J, Lai B, Qin Y, Wang H, Hu G. LcGST4 is an anthocyanin-related glutathione S-transferase gene in Litchi chinensis Sonn. Plant Cell Rep 2016;35:831–843.

31 Boss PK, Davies C, Robinson SP. Analysis of the expression of anthocyanin pathway genes in developing Vitis vinifera L. cv Shiraz grape berries and the implications for pathway regulation. Plant Physiol 1996;111:1059–1066.

32 Jaakola L. New insights into the regulation of anthocyanin biosynthesis in fruits. Trends Plant Sci 2013;18:477–483.

33 Montefiori M, Espley RV, Stevenson D, et al. Identification and characterisation of F3GT1 and F3GGT1, two glycosyltransferases responsible for anthocyanin biosynthesis in red-fleshed kiwifruit (Actinidia chinensis). The Plant Journal 2011;65:106–118.

34 Roldan MVG, Outchkourov N, van Houwelingen A, et al. An O-methyltransferase modifies accumulation of methylated anthocyanins in seedlings of tomato. Plant J 2014;80:695–708.

35 Zhao X, Yuan Z, Yin Y, Feng L. Patterns of pigment changes in pomegranate (Punica granatum L.) peel during fruit ripening. Acta Hortic 2015;1089:83–89.

36 Xu Q, Chen L-L, Ruan X, et al. The draft genome of sweet orange (Citrus sinensis). Nat Genet 2013;45:59–66.

37 Seitz C, Ameres S, Schlangen K, Forkmann G, Halbwirth H. Multiple evolution of flavonoid 3′,5′-hydroxylase. Planta 2015;242:561–573.

38 Wang YS, Xu YJ, Gao LP, et al. Functional analysis of favonoid 3′,5′-hydroxylase from Tea plant (Camellia sinensis): critical role in the accumulation of catechins. BMC Plant Biol 2014;14:347.

39 Petroni K, Tonelli C. Recent advances on the regulation of anthocyanin synthesis in reproductive organs. Plant Sci 2011;181:219–229.

40 Zhou H, Lin-Wang K, Wang H, et al. Molecular genetics of blood-fleshed peach reveals activation of anthocyanin biosynthesis by NAC transcription factors. The Plant Journal 2015;82:105–121.

41 Dahlgren R, Thorne RF. The order Myrtales: circumscription, variation, and relationships. Ann Mo Bot Gard 1984;71:633–699.

42 Schaefer H, Renner S Cucurbitaceae. In: Kubitzki K, eds. The families and genera of vascular plants. Berlin: Springer, 2011: 112–174.

43 Zhang ZY, Lu AM, William GDA Solanaceae. In: Wu ZY, Raven PH, Hong DY, eds. Flora of China. Beijing: Garden Press, 1994.

44 Chan AP, Crabtree J, Zhao Q, et al. Draft genome sequence of the oilseed species Ricinus communis. Nat Biotechnol 2010;28:951–U953.

45 Brambilla V, Battaglia R, Colombo M, et al. Genetic and molecular interactions between BELL1 and MADS box factors support ovule development in Arabidopsis. Plant Cell 2007;19:2544–2556.

46 Duszynska D, McKeown PC, Juenger TE, Pietraszewska-Bogiel A, Geelen D, Spillane C. Gamete fertility and ovule number variation in selfed reciprocal F1 hybrid triploid plants are heritable and display epigenetic parent-of-origin effects. New Phytol 2013;198:71–81.

47 Narzary D, Rana TS, Ranade SA. Genetic diversity in inter-simple sequence repeat profiles across natural populations of Indian pomegranate (Punica granatum L.). Plant Biol 2010;12:806–813.

48 O′Connell J, Schulz-Trieglaff O, Carlson E, Hims MM, Gormley NA, Cox AJ. NxTrim: optimized trimming of Illumina mate pair reads. Bioinformatics 2015;31:2035–2037.

49 Butler J, MacCallum I, Kleber M, et al. ALLPATHS: De novo assembly of whole-genome shotgun microreads. Genome Res 2008;18:810–820.

50 Boetzer M, Henkel CV, Jansen HJ, Butler D, Pirovano W. Scaffolding pre-assembled contigs using SSPACE. Bioinformatics 2011;27:578–579.

51 Luo R, Liu B, Xie Y, et al. SOAPdenovo2: an empirically improved memory-efficient short-read de novo assembler. Gigascience 2012;1:18.

52 Xu Z, Wang H. LTR_FINDER: an efficient tool for the prediction of full-length LTR retrotransposons. Nucleic Acids Res 2007;35:W265–W268.

53 Han Y, Wessler SR. MITE-Hunter: a program for discovering miniature inverted-repeat transposable elements from genomic sequences. Nucleic Acids Res 2010;38:e199.

54 Price AL, Jones NC, Pevzner PA. De novo identification of repeat families in large genomes. Bioinformatics 2005;21 Suppl 1:i351–358.

55 Edgar RC, Myers EW. PILER: identification and classification of genomic repeats. Bioinformatics 2005;21 Suppl 1:i152–158.

56 Wicker T, Sabot F, Hua-Van A, et al. A unified classification system for eukaryotic transposable elements. Nat Rev Genet 2007;8:973–982.

57 Bao W, Kojima KK, Kohany O. Repbase Update, a database of repetitive elements in eukaryotic genomes. Mob DNA 2015;6:11.

58 Haas BJ, Papanicolaou A, Yassour M, et al. De novo transcript sequence reconstruction from RNA-Seq using the Trinity platform for reference generation and analysis. Nat Protoc 2013;8:1494–1512.

59 Stanke M, Keller O, Gunduz I, Hayes A, Waack S, Morgenstern B. AUGUSTUS: ab initio prediction of alternative transcripts. Nucleic Acids Res 2006;34:W435–W439.

60 Campbell MA, Haas BJ, Hamilton JP, Mount SM, Buell CR. Comprehensive analysis of alternative splicing in rice and comparative analyses with Arabidopsis. BMC Genomics 2006;7:327.

61 Korf I. Gene finding in novel genomes. BMC Bioinformatics 2004;5:59.

62 Majoros WH, Pertea M, Salzberg SL. TigrScan and GlimmerHMM: two open source ab initio eukaryotic gene-finders. Bioinformatics 2004;20:2878–2879.

63 Burge C, Karlin S. Prediction of complete gene structures in human genomic DNA. J Mol Biol 1997;268:78–94.

64 Parra G, Blanco E, Guigó R. GeneID in Drosophila. Genome Res 2000;10:511–515.

65 Schneider M, Tognolli M, Bairoch A. The Swiss-Prot protein knowledgebase and ExPASy: providing the plant community with high quality proteomic data and tools. Plant Physiol Biochem 2004;42:1013–1021.

66 Birney E, Clamp M, Durbin R. GeneWise and Genomewise. Genome Res 2004;14:988–995.

67 Wu TD, Watanabe CK. GMAP: a genomic mapping and alignment program for mRNA and EST sequences. Bioinformatics 2005;21:1859–1875.

68 Elsik CG, Mackey AJ, Reese JT, Milshina NV, Roos DS, Weinstock GM. Creating a honey bee consensus gene set. Genome Biol 2007;8:R13.

69 Wang Y, Tang H, DeBarry JD, et al. MCScanX: a toolkit for detection and evolutionary analysis of gene synteny and collinearity. Nucleic Acids Res 2012;40:e49.

70 Edgar RC. MUSCLE: multiple sequence alignment with high accuracy and high throughput. Nucleic Acids Res 2004;32:1792–1797.

71 Hasegawa M, Kishino H, Yano T-A. Dating of the human-ape splitting by a molecular clock of mitochondrial DNA. J Mol Evol 1985;22:160–174.

72 Li L, Stoeckert CJ, Roos DS. OrthoMCL: identification of ortholog groups for eukaryotic genomes. Genome Res 2003;13:2178–2189.

73 Guindon S, Dufayard J-F, Lefort V, Anisimova M, Hordijk W, Gascuel O. New algorithms and methods to estimate maximum-likelihood phylogenies: Assessing the performance of PhyML 3.0. Syst Biol 2010;59:307–321.

74 Hedges SB, Dudley J, Kumar S. TimeTree: a public knowledge-base of divergence times among organisms. Bioinformatics 2006;22:2971–2972.

75 Albertin CB, Simakov O, Mitros T, et al. The octopus genome and the evolution of cephalopod neural and morphological novelties. Nature 2015;524:220–224.

76 Besemer J, Lomsadze A, Borodovsky M. GeneMarkS: a self-training method for prediction of gene starts in microbial genomes. Implications for finding sequence motifs in regulatory regions. Nucleic Acids Res 2001;29:2607–2618.

77 Trapnell C, Roberts A, Goff L, et al. Differential gene and transcript expression analysis of RNA-Seq experiments with TopHat and Cufflinks. Nat Protoc 2012;7:562–578.

78 Anders S, Huber W. Differential expression analysis for sequence count data. Genome Biol 2010;11:R106.

79 Lan P, Li W, Wen T-N, et al. iTRAQ Protein Profile Analysis of Arabidopsis Roots Reveals New Aspects Critical for Iron Homeostasis. Plant Physiol 2011;155:821–834.

